# Older Women with lower lean mass values have hypermethylated sites in the PI3K-Akt pathway

**DOI:** 10.1101/2022.10.14.512202

**Authors:** Igor Massari Correia, Guilherme da Silva Rodrigues, Natália Yumi Noronha, Mariana Luciano de Almeida, Andressa Crystine da Silva Sobrinho, Carla Barbosa Nonino, Carlos Roberto Bueno Júnior

## Abstract

The increase in lean mass is directly related to the loss of independence, muscle strength, and worse quality of life over the years. Studies in epigenetics can provide accurate answers about lean mass, demonstrating changes in DNA methylation patterns and possible changes in gene expression. The objective of this study was to verify whether there is a difference in the methylation profile among Brazilian women aged 50 to 70 years with greater or lesser lean mass. A cross-sectional study comprised 22 women aged 50 to 70 years, with 2 groups of 11 participants (Low Lean Mass and More Lean Mass). Lean mass was measured by dual-energy X-ray emission densitometry (DEXA). Blood DNA was collected for methylation assays using the Illumina 850k EPIC Infinium Methylation BeadChip, analyzing data from the Bioconductor chAMP data package medium in RStudio software. We obtained 1,913 differentially methylated (p ≤ 0.005 of delta β > 5% and delta β < −5 %) with a total of 979 genes with different methylation sites between groups (p ≤ 0.005; −5% > delta β > 5%). In addition, the pathway with the greatest power of significance was PI3K-Akt, presenting an FDR of 4.6 x 10^-3^. Thus, our results demonstrate a differentiation between specific sites of different genes, which have essential functions in body composition and energy metabolism, supporting future studies that aim to relate lean mass with epigenetics.

## INTRODUCTION

Age is one of the main predictors of chronic diseases, which are related to the majority of morbidities, hospitalizations, health costs, and mortality, representing a risk factor for geriatric syndromes, frailty, immobility, and decreased physical resilience. In parallel, in the aging process, there is a decrease in cell regeneration capacity and a simultaneous increase in cell apoptosis, which is related to the loss of cardiorespiratory function and the decrease in muscle mass and strength. Lean mass plays an important role in the aging process, as its decrease is directly associated with loss of independence, muscle strength, and lower quality of life over the years (1). The lean mass measured by dual-energy X-ray emission densitometry (DEXA) encompasses muscles, organs, and body fluids, disregarding fat mass and bone mass.

Studies show that low muscle mass is a predictor of mortality in older adults - reduced levels of muscle mass can increase the risk of mortality in this population by 11 times, and when associated with death from cardiovascular diseases, the risk of mortality increases by 14 times (2). Around age 40, a more pronounced decrease in lean mass can be observed, which can result in sarcopenia, which is defined as the involuntary loss of skeletal muscle mass and a decrease in strength related to age (3). This loss can be greater or lesser depending on the lifestyle of each individual, especially regarding the practice of physical activities and diet. That is, people who eat properly and are physically active are less likely to develop sarcopenia (4). Sarcopenia can be considered a health problem due to its aggravating consequences on muscle functionality and body composition, and studies that focus on its prevention are lacking. Currently, the annual health costs related to lean mass reduction in the United States are estimated at more than 18 billion dollars, as lean mass reduction is associated with diseases such as obesity, diabetes, and osteoporosis (5). In Brazil, the costs of the Unified Health System (SUS) were 3.45 billion reais in 2018 (more than 890 million dollars) - 72% of the costs were related to individuals between 30 and 69 years of age and 56% to women. According to the analysis of the World Economic Bank, it is estimated that countries such as Brazil, China, India, and Russia lose more than 20 million productive years of life per year as a result of chronic non-communicable diseases (NCDs), generating a loss in the Brazilian economy of US$ 4.18 billion between 2006 and 2015.

Studies in epigenetics can provide precise answers regarding physiological responses at the molecular level and the modification of lean body mass in the human body since, with the aging process, there is a decrease in this body tissue and a change in the DNA methylation pattern, making possible the modification of gene expression related to the regulation of lean mass (6). Hypermethylation, in most cases, leads to gene silencing, while hypomethylation leads to increased gene expression (7,8). In this way, knowing the methylation profile according to lean mass could be a factor that will help in decision-making by health professionals in preventive measures against the reduction in this body tissue and physiological alterations related to this reduction.

There is evidence in the literature suggesting the influence of DNA methylation on the expression of metabolically essential genes in skeletal muscle and its possible implication in susceptibility to age-related metabolic diseases (9). However, little is known about methylation patterns in relation to lean mass in healthy older people, and more evidence is required in this population. (9). One study compared the DNA methylation profile in the skeletal muscle of 24 older males with that of 24 young males of the same sex. The authors identified 5,963 differentially methylated CpG sites between the two groups, and most of these (92%) were hypermethylated in the older group, suggesting epigenetic links between post-mitotic skeletal muscle aging and DNA methylation (10). In a longitudinal study carried out with 1550 middle-aged female twins, the authors identified seven regions whose methylation status was significantly associated with the variation in skeletal muscle mass, demonstrating that epigenetic studies have an essential role in the study of muscle mass (11).

It is important to emphasize that there is no study comparing the DNA methylation profile of older people with different values of lean mass. Thus, the objective of the present study was to look for differentially methylated sites and establish whether there is any relationship between the difference in lean mass between a group that has a higher percentage of lean mass and a group that has a lower percentage of lean mass in Brazilian women aged 50 to 70 years old. Our hypothesis is that females with a higher percentage of lean mass have a different methylation profile from females with a lower percentage of lean mass.

## METHODS

The research project was submitted and approved by the Ethics Committee for Research with Human Beings of EEFERP-USP (CAAE: 79582817.0.0000.5659), based on Resolution 466/12 of the Ministry of Health, regulated by the National Health Council. This study was registered in the Brazilian Registry of Clinical Trials (REBEC) with the registration number RBR-3g38dx.

Women aged 50 to 70 years enrolled in the Physical Education Program for the Elderly (PEFI) of the School of Physical Education and Sport at USP de Ribeirão Preto (EEFERP-USP) were invited to participate in our study. In total, 40 women made up the sample of this study. After measuring lean mass by DEXA, these participants were divided into two groups: *more lean mass* (MLM) and *less lean mass*(LLM). The MLM group included the 20 participants with the highest percentage of lean mass, and the LLM group included the 20 participants with the lowest percentage of lean mass, measured by DEXA.

### Physical and functional assessment instruments

Anthropometric assessments were used to characterize the participants: body mass, height, body mass index (BMI), and waist circumference (WC) (12). The Food Consumption Markers Form (FMCA) of the Ministry of Health was used to assess the participants’ diet, and the modified Baecke Questionnaire for the elderly to assess their physical activity level (MBQE) (13,14).

The following motor tests from the Rikli and Jones battery of tests were used: elbow flexion and extension (EFC) test to assess upper limb strength, sit-to-stand test (STS) to assess lower limb strength, and 6-minute walk (6MWT) to assess the participants’ aerobic fitness (15). In addition, we performed the maximum repetitions test to complement the participants’ strength assessment, using the incline bench press and the 45° leg press for this purpose (16).

### DEXA Evaluation

A laboratory technician duly trained for these assessments performed measurement and body composition analyses following standard procedures. At the time of the evaluations, the participants were not wearing any accessories, such as necklaces, bracelets, and earrings, to reduce errors in the results (17,18).

The whole-body scans were manually analyzed for total and percentage lean tissue mass and total and percentage fat mass (17,18).

### Blood tests

Blood tests for glucose, insulin, and glycated hemoglobin were performed after the participants had fasted for 12 hours. The blood material was immediately sent to the Faculdade de Ciências Farmacêuticas da USP de Ribeirão Preto (FCFRP), wrapped in ice in a thermal box in order to preserve the material, and analyzed using the *Wiener Lab BT 3000* plus autoanalyzer (19). The reagents used for the analyses described in this topic were from the same batch (LABORLAB).

### DNA extraction and methylation assay

A peripheral blood sample was collected in tubes with EDTA in a sterile environment, and DNA was extracted from leukocytes. Genomic DNA was isolated from 500 μL of peripheral blood by the *salting out* method (20). DNA concentration and quality were evaluated by spectrophotometry at 260 and 280 nm (A260 and A280) using Biodrop, and integrity was evaluated using 1% agarose gel - the run protocol adopted was 90 minutes at 80 W (21). In addition, the extraction technique was evaluated using negative controls. Subsequently, the samples were stored at −20 °C. Leukocyte genomic DNA (500 ng) was treated with bisulfite using the DNA EZ Methylation-Gold methylation kit (Zymo Research, CA, USA), following the recommendations stipulated by the manufacturer.

DNA methylation assays were performed using the *array* technique, with the Illumina EPIC *Infinium Methylation BeadChip*, following the manufacturer’s instructions (22,23)*. The Illumina iScan system detects Infinium BeadChip* fluorescence, the intensities are exported as *raw data* using the chAMP package (for the R platform), and the data are converted into beta methylation values, processed, and analyzed (24,25).

### DNA methylation analysis

Methylation analyses were performed using RStudio software and calculated using the Bioconductor chAMP data package. The DMP function based on the limma algorithm was used to find the differentially methylated sites between the two groups, classified as hypermethylated and hypomethylated, which presented values of p ≤ 0.005 and delta β > 5% for hypermethylated sites and p ≤ 0.005 and delta β < −5% for hypomethylated sites. We also adopted the DMR function from the same package, which presents the differentially methylated regions, to analyze whether, in addition to DMPs, the difference in lean mass is also reflected in the differentially methylated regions.

### Enrichment analysis

Enrichment was obtained through gene enrichment analysis (GSEA). This method is based on comparing a list of genes (LG), ordered according to their correlation with the phenotype, and the genes described in a specific metabolic pathway (MP). To this end, calculations of the enrichment index, significance levels, and corrected significance levels for multiple tests are performed. The GSEA evaluates each gene independently, ignoring the relationships between them. The ShinyGO v0.66: Gene Ontology Enrichment Analysis database was used to perform interaction and functional enrichment. We adopted an FDR < 0.05 for the human organism to select the gene-related KEGG metabolic pathways to analyze the results (26).

### Statistical analysis

Two researchers entered the characterization data of the participants and biochemical variables into Excel spreadsheets, and then the information was cross-referenced to eliminate possible typing errors. We used the Statistical Package for the Social Sciences (SPSS) software for statistical processing. We used the Shapiro Wilk as a normality test, and as the data presented parametric distribution, we adopted the Student-t test, p < 0.05.

The M value was adopted to compare the difference in methylation of the sites of the genes that showed statistical differences by DMR, which were selected and passed through the quality control of chAMP and Bumpunter (p ≤ 0.05). In order to use the M value, it was necessary to transform the beta values into the M value. This approach has been widely used in the analysis of DNA methylation microarrays, as it increases the range of values, thus better performing statistical analyses than the beta values. The beta values (*Beta*) of the sites were converted into M values (*M_i_*) using the following equation on a logarithmic scale (24,27):

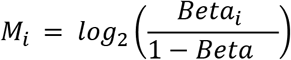

## RESULTS

There was a similarity between the groups in terms of age, motor, and functional tests (RM, STS, EFC, and 6MWT), waist circumference, and assessments of the participant’s level of physical activity and daily diet (MBQE and FMCA) (Table 1). Body composition measurements were the only variables that showed significant differences between groups (BMI and lean mass percentage - p < 0.05). In table 1 it is possible to observe this difference for the percentage of lean mass (MLM: 56.7 ± 2.8%; LLM 49.1 ± 2.5%) and for BMI (MLM: 27.7 ± 3.4 kg.m^-2^; LLM: 31.7 ± 3.6 kg.m^-2^). Although participants in the MLM group presented significantly greater lean mass than participants in the LLM group (p < 0.001), the groups did not show differences in performance in functional tests.

**Table 1.** Anthropometric data and physical fitness profiles in women aged 50 to 70 years, separated by groups (LLM and MLM).

|  | Less Lean Mass<br>(n=20) | More Lean Mass<br>(n=20) | p-Value |
| --- | --- | --- | --- |
| Lean Mass (%) | 49.1 ± 2.5 | 56.7 ± 2.8* | < 0.001 |
| Age (years) | 61.1 ± 5.3 | 60.5 ± 4.2 | 0.626 |
| BMI (kg/m<sup>2</sup>) | 31.7 ± 3.6 | 27.7 ± 3.4* | 0.001 |
| WC (cm) | 96.9 ± 8.0 | 91.9 ± 12.2 | 0.140 |
| EFC (reps) | 14.9 ± 5.0 | 17.3 ± 4.3 | 0.103 |
| RM supine (kg) | 22.4 ± 4.4 | 22.1 ± 4.2 | 0.871 |
| RM leg press (kg) | 130.7 ± 28.4 | 123.9 ± 40.8 | 0.558 |
| STS (reps) | 22.1 ± 8.1 | 18.3 ± 8.0 | 0.152 |
| 6MWT (m) | 538.2 ± 64.7 | 523.8 ± 59.3 | 0.466 |
| MBQE (points) | 5.8 ± 3.8 | 4.0 ± 2.8 | 0.106 |
| FMCA (points) | 17.3 ± 10.3 | 19.6 ± 8.7 | 0.473 |

Legend - BMI: body mass index; WC: waist circumference; EFC: elbow flexion and extension; RM: maximum repetition; SEL: sit and stand; 6MWT: 6-minute walk test; BME: Modified Baecke Questionnaire for the Elderly; FMCA: food consumption markers form; cm: centimeters; m: meters; reps: repetitions; kg: kilos; m: meters. \*Student-t test, $p < 0.05$ .

After quality control performed with the chAMP package, we obtained 719,902 valid sites - of these 9,032 were significantly different between groups (p ≤ 0.005) and 1,913 were differentially methylated (p ≤ 0.005 of delta β > 5% and delta β < −5 %). Figure 1A shows the differentially hypermethylated sites in the LLM group compared to the MLM group, totaling 1,776 sites, distributed in 1stExon, 3’UTR, 5’UTR, Body, ExonBnd, TSS1500, TSS200, and IGR. It is possible to notice that the regions of the open sea CpG islands, together with the regions of the Body and IGR genes, presented the highest frequency of hypermethylation in the LLM group.

**Figure 1A.**
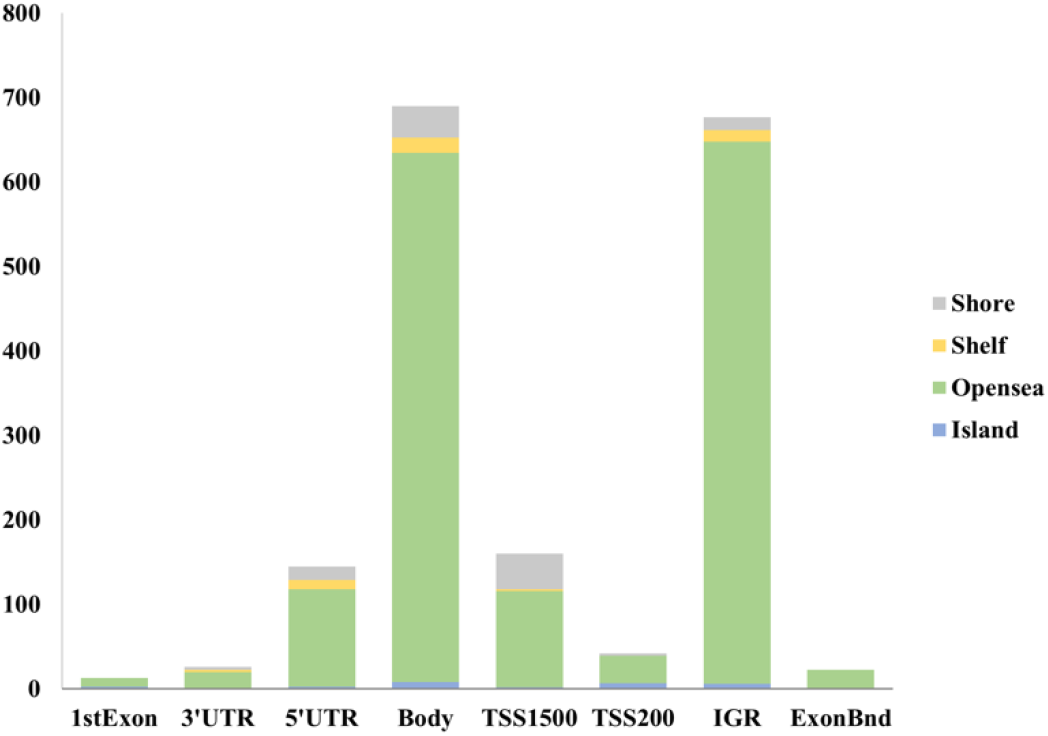
The number of hypermethylated sites per gene region in the LLM group (1,776). The X-axis of the graph shows the regions of genes and intergenic regions, and the Y-axis shows the number of hypermethylated sites. Each color represents a region of the CpG islands related to hypermethylated genes.

In Fig. 1B, the hypomethylated sites in the LLM group are shown in relation to the MLM group - 137 sites are distributed in the regions of the genes (a large proportion in the body and the IGR of the genes) so that the open sea and shore CpG regions were the ones that showed the highest frequency of hypomethylation.

**Figure 1B.**
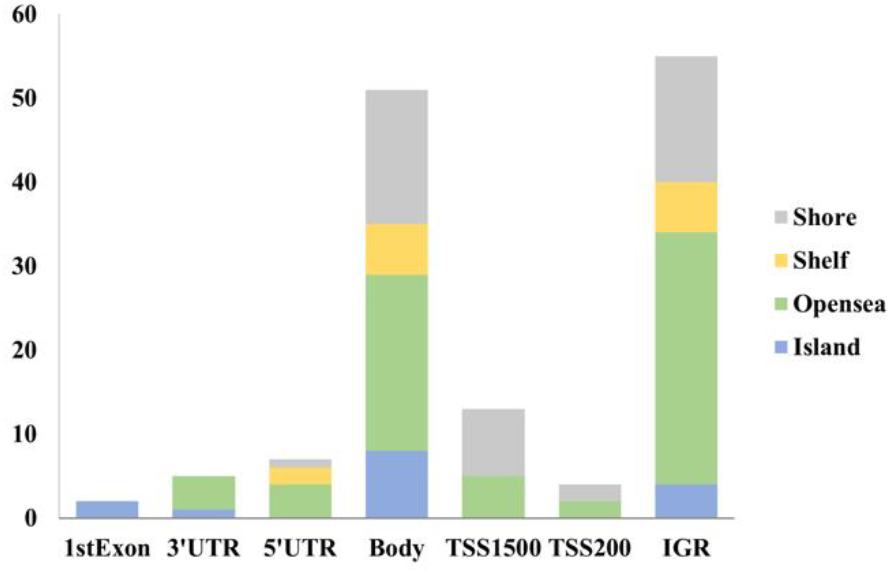
The number of hypomethylated sites per gene region in the LLM group (137). The X-axis of the graph shows the regions of genes and intergenic regions, and the Y-axis is the number of hypomethylated sites. Each color represents a region of the CpG islands related to hypomethylated genes.

The results showed a total of 979 genes with differently methylated sites between groups (p ≤ 0.005; −5% > delta β > 5%). The sites were differently hypermethylated in 897 and differently hypomethylated in 82 in the LLM group. Figure 2A represents the enrichment of the genes that had hypermethylated sites in the LLM group, demonstrating the FDR values of KEGG pathways (< 0.05) and the number of these genes related to these pathways. However, we could not find significant relationships for the enrichment of genes with hypomethylated sites in the LLM group.

**Figure 2A.**
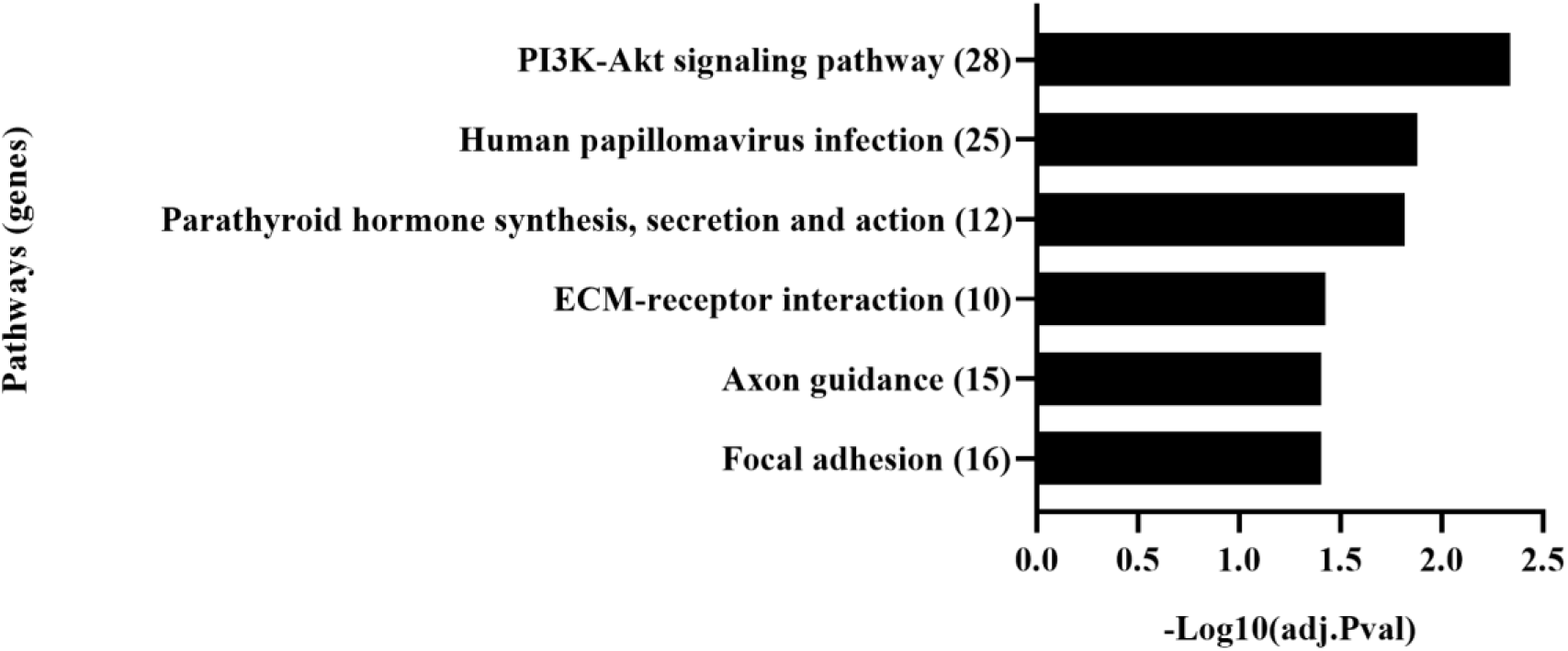
KEGG pathway enrichment analysis of hypermethylated genes in the LLM group. The names of KEGG pathways are present on the vertical axis with the number of genes related to them in parentheses. The horizontal axis represents the FDR values in −Log of 10. For this analysis, the homo sapiens reference organism was used, considering an FDR < 0.05.

The KEGG pathways tree in Fig. 2B demonstrates the connections between the pathways and the absolute value of the FDR of each one. Thus, it is possible to notice that there is an interaction between these pathways, so that the pathway with the greatest power of significance was PI3K-Akt, presenting an FDR of 4.6 x 10^-3^, which is related to the metabolism of lean mass, as can be seen in Fig. 3, demonstrating the signals and actions in the human body.

**Figure 2B.**
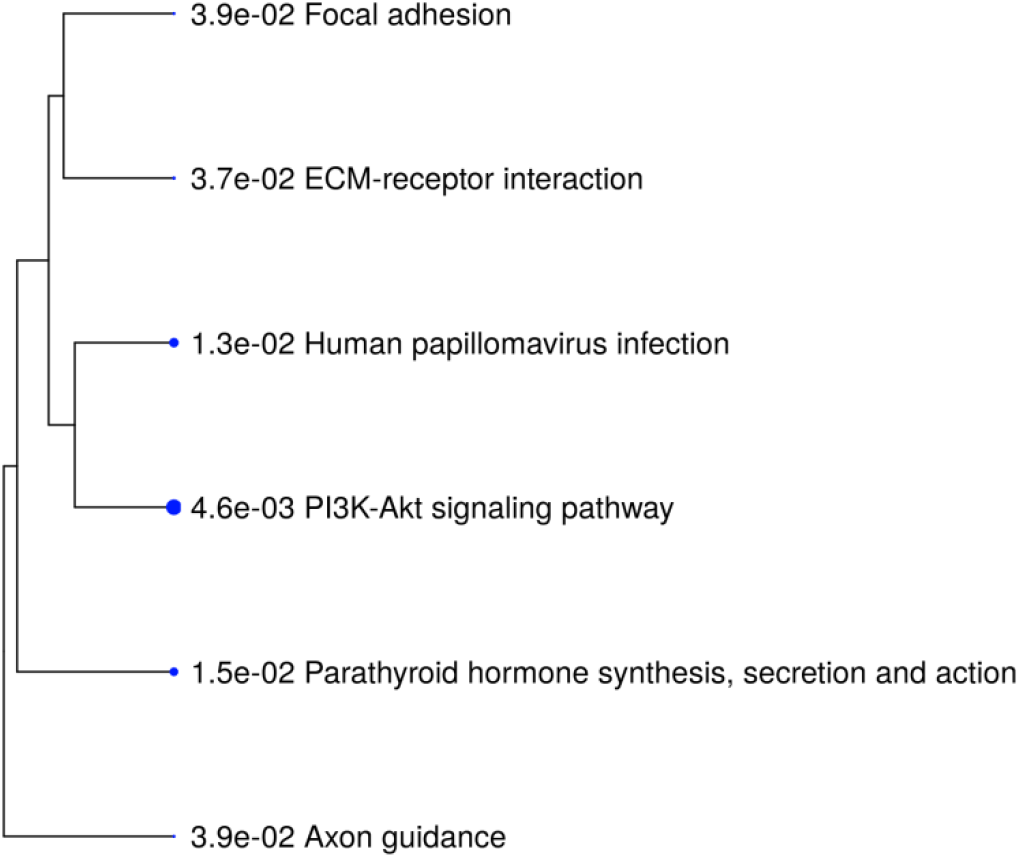
Enrichment analysis tree of hypermethylated genes in the LLM group. The hierarchical clustering tree summarizes the correlation between the significant pathways listed in Fig. 2A. Pathways with many shared genes were grouped together and larger dots indicate more significant FDR values.

**Figure 3.**
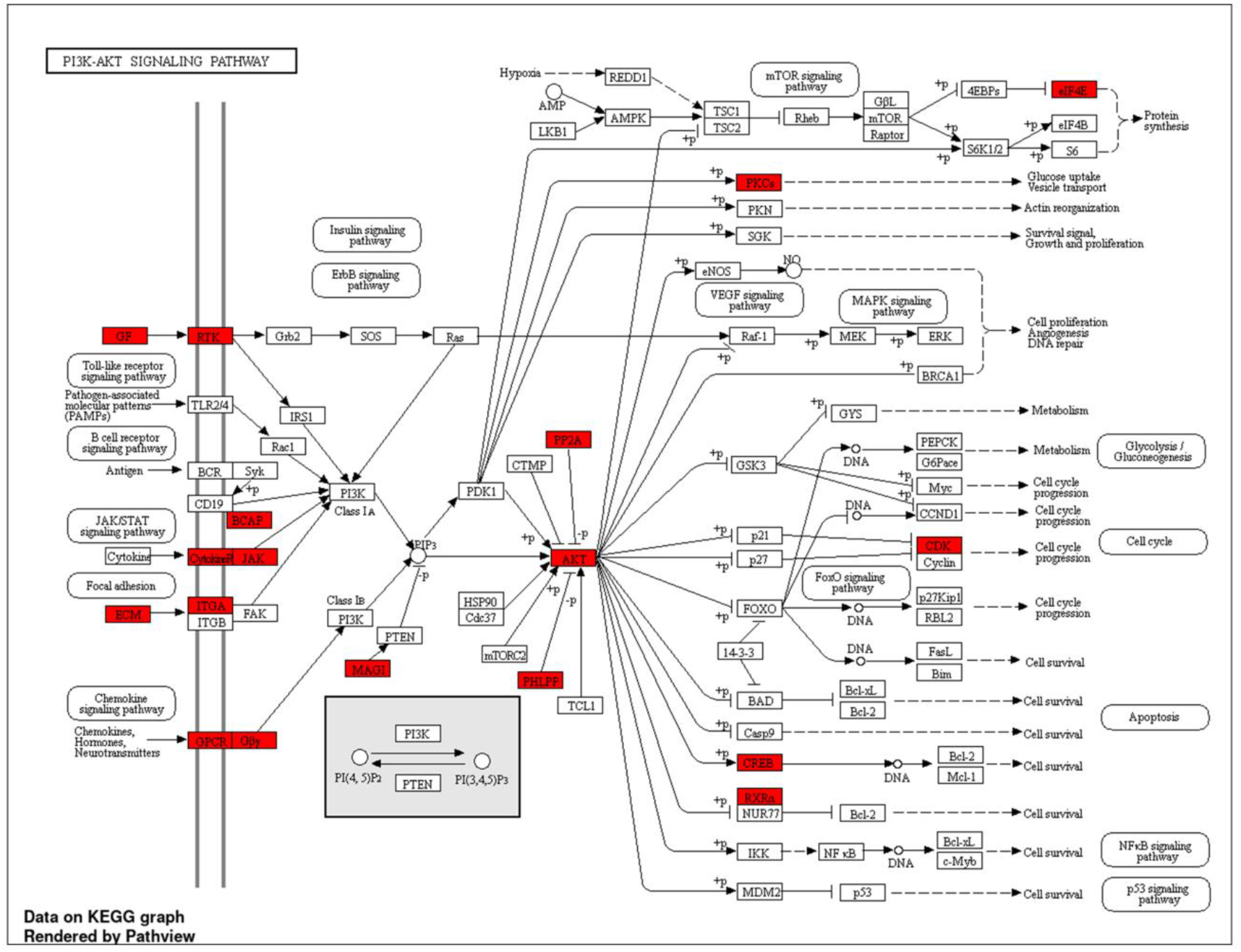
PI3K-Akt KEGG pathway. This image represents the signals and actions of the PI3K-Akt pathway in the human body. The genes highlighted in red showed differences in the methylation profile between the groups.

The analysis of blood tests became relevant following the result of the enrichment of genes that were differently methylated between the MLM and LLM groups, since the PI3K-Akt pathway showed a greater significant difference and is also related to the glucose pathway in the human body. (Fig. 3). Thus, Table 2 presents an analysis of the glucose and insulin profile in the participants’ blood. However, it was not possible to observe a significant difference (p < 0.05) between the groups in the results of blood tests, which strengthens the hypothesis that changes in this intracellular signaling pathway are associated with muscle trophism.

**Table 2.**
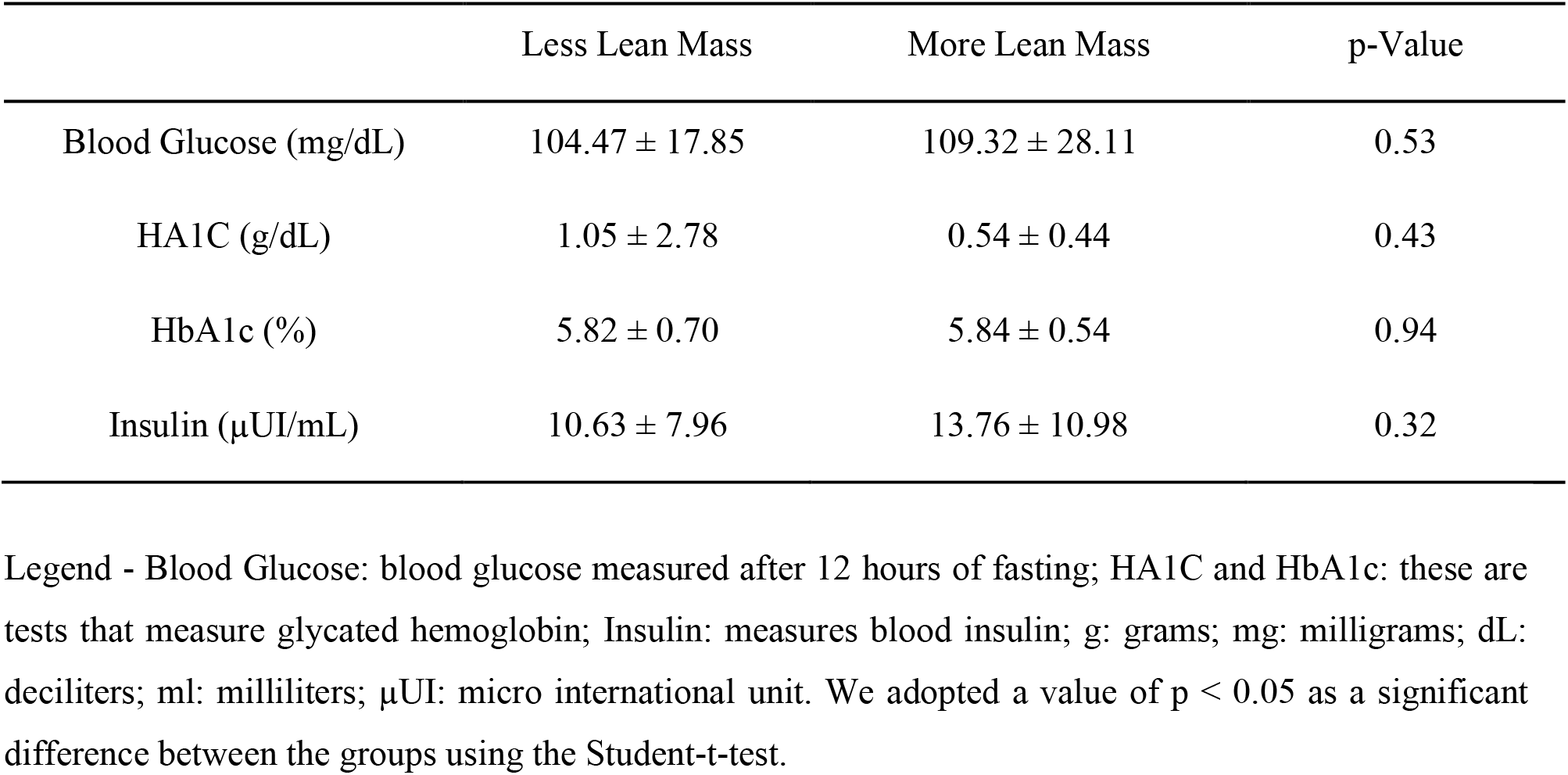
Blood glucose and insulin profile in women aged 50 to 70 years separated by groups (LLM and MLM).

|  | Less Lean Mass | More Lean Mass | p-Value |
| --- | --- | --- | --- |
| Blood Glucose (mg/dL) | 104.47 ± 17.85 | 109.32 ± 28.11 | 0.53 |
| HA1C (g/dL) | 1.05 ± 2.78 | 0.54 ± 0.44 | 0.43 |
| HbA1c (%) | 5.82 ± 0.70 | 5.84 ± 0.54 | 0.94 |
| Insulin (μUI/mL) | 10.63 ± 7.96 | 13.76 ± 10.98 | 0.32 |

Legend - Blood Glucose: blood glucose measured after 12 hours of fasting; HA1C and HbA1c: these are tests that measure glycated hemoglobin; Insulin: measures blood insulin; g: grams; mg: milligrams; dL: deciliters; ml: milliliters; $\mu$ UI: micro international unit. We adopted a value of $p < 0.05$ as a significant difference between the groups using the Student-t-test.

DMR analysis revealed that the Bumphunter detected 10 DMRs with a P value ≤ 0.05. Of the 10 DMRs, one was in the intergenic region, and the other nine were linked to genes. The beta values were transformed into M values, and the sites that showed statistically significant differences between the groups are shown in Fig. 4. It is possible to notice a pattern in the different levels of methylation in the comparisons between groups. The MLM group was shown to be hypermethylated about the LLM group in the genes in Fig 4. Notably, only the C170rf97 gene was in the TSS1500 promoter zone, and the others were in the body region. The HLA-DPB15 and C170rf97 genes showed differentially methylated sites in the CpG island region, while the others were in the shore region. The M values of the other sites, respective genes, and methylation types are presented in Supplementary file 1.

**Figure 4.**
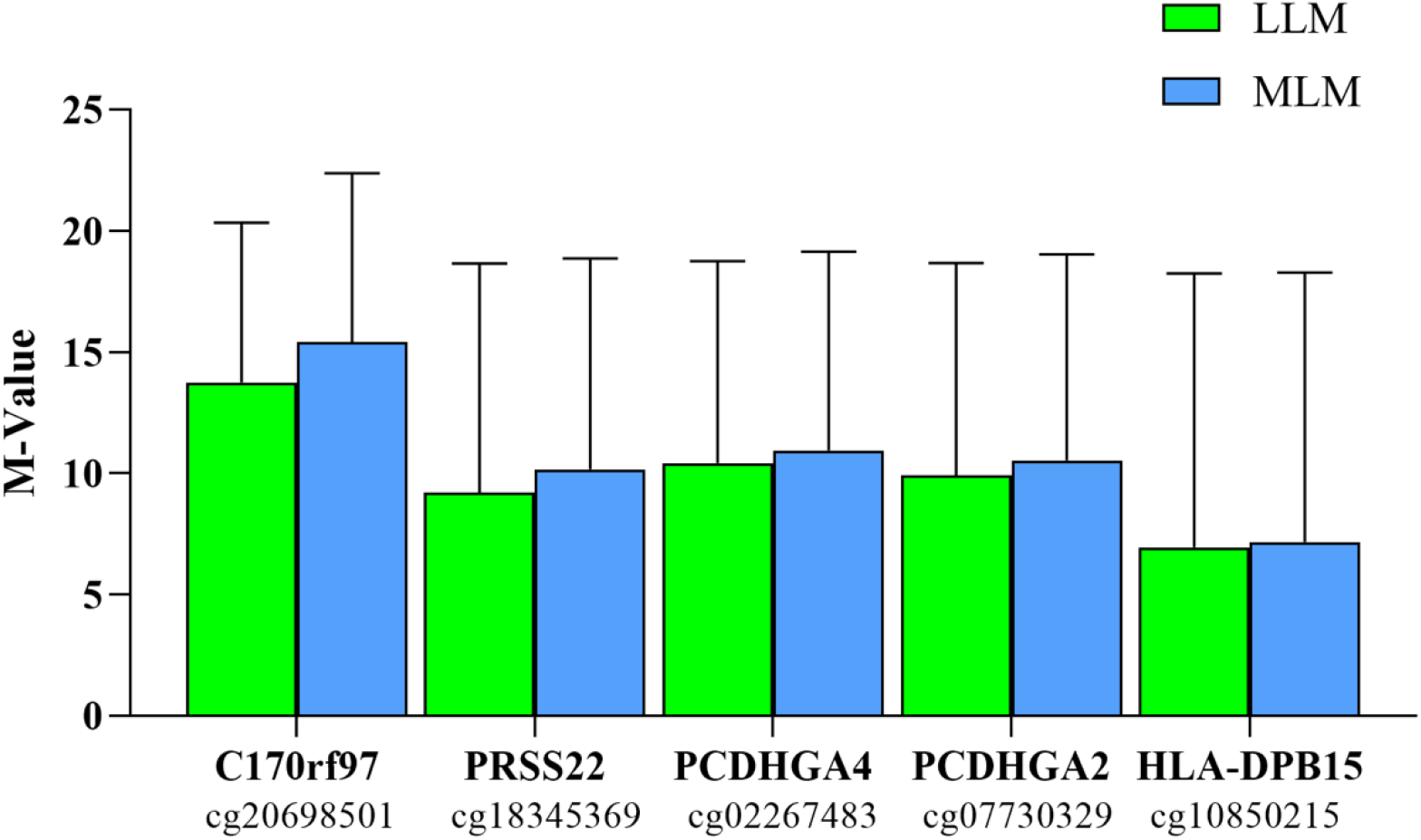
M-value of differently methylated sites between groups and their respective genes derived from DMR analysis. The gene names are on the X-axis, and the identification of the sites of each gene is just below them. The sites that were inserted in this group are methylated differently between the groups (p ≤ 0.05).

## DISCUSSION

The current study aimed to analyze the differences in the methylation profile between older women with a higher percentage of lean mass and those with a lower percentage of lean mass. We observed a significant difference in BMI between the groups. Participants with higher lean mass presented lower BMI values, indicating lower body fat mass. This difference in body composition between the LLM and MLM groups was related to the difference in the methylation profile between these groups, mainly in the PI3-Akt pathway.

Increased body fat, decreased strength, and lean mass increase mortality risks (2,28). The lean mass measured by DEXA is made up of muscles, organs, and body fluids, excluding fat mass. Therefore, a participant in the MLM group who has the same weight as a participant in the LLM group presents better health indicators, as she has a lower percentage of fat mass and a higher percentage of lean mass in her body composition, which may be related to a lower risk of mortality compared to the participant in the LLM group. One of the explanations for the differences in body composition between individuals is that there is a link between genetics, environment, and lifestyle, which can facilitate or hinder the improvement of these health indicators (29). A study with mice showed that the inactivation of the melanocortin-3 receptor caused an increase in body fat and a reduction in lean mass (30). In this way, it is possible to observe that some genes can contribute to the modification of body composition according to their expression, which is essential for lifestyle interventions, as gene expression is more related to lifestyle than genetics alone (29).

Our study demonstrated a significant difference in the methylation profile for the genes of the PI3K/Akt pathways, human papillomavirus (HPV) infection, parathyroid hormone, extracellular matrix (ECM) receptor interaction, axon orientation, and focal adhesion. The genes related to these pathways had their sites hypermethylated in the LLM group compared to the MLM group. These results indicate a possible difference in gene expression between the groups (7,8).

HPV is composed of tumorigenic DNA that infects the basal cells of the epithelium - it can follow several routes to enter the cells. The expression of genes in this pathway is related to positive regulation of the PI3K pathway, which does not depend on the Akt protein to continue functioning. However, HPV impairs the activation of the Akt protein, decreasing the action of the protein and the signaling related to the integral functioning of the PI3K/Akt pathway (31–33).

The parathyroid hormone pathway is related to bone remodeling and the regulation of calcium homeostasis (34). In addition, the parathyroid hormone-related protein positively regulates the expression of α 6 β 4 integrin, favoring the functioning of the PI3K/Akt pathway. This relationship results in the suppression of cellular apoptosis (35).

The ECM interaction pathway is essential for regulating intercellular interaction and tissue architecture. The ECM undergoes considerable remodeling in situations of injury, repair, short-term overeating, metabolic dysfunction, and obesity (36–39). Studies demonstrate that the ECM pathway can increase biochemical signals, stimulating myogenesis. In addition, it is related to the regulation of insulin signaling in cells (36–39). When insulin is bound to cell membrane receptors, signal transduction pathways activate PI3K-Akt and insulin receptor substrate (IRS) −1, which act on glucose uptake through GLUT4. Therefore, low expression of the genes of the ECM interaction pathway can result in impairment in poststimulus insulin signaling, which can lead to the development of insulin resistance and type 2 diabetes mellitus (36–39).

The axon guidance (OA) pathway is related to the connectivity and repair of neurons throughout life and guides the brain’s wiring during fetal development. The neuron signaling pathway complex and its intracellular signaling are related to various cellular responses: trafficking events, gene transcription, cytoskeletal remodeling, ubiquitination, and protein translation (40–42). The PI3K pathway assists the OA pathway in controlling neuronal migration, neuronal morphogenesis, and the development of dendrites and synapses. These pathways are essential in stages of brain development, which include mediating extrinsic and intrinsic signals, such as growth factors, axon guidance, extracellular matrix components, and signaling components that act in translation. However, the literature does not explicitly state a direct relationship between this pathway and lean mass or glucose metabolism (40,41).

Focal adhesions (FA) are sites where there is adhesion mediated by integrin and proteoglycan bound to the actin cytoskeleton. These sites are modified by cells in response to alterations in molecular composition, structure, and physical forces present in the extracellular matrix. FA regulates integrin signaling and function complexes, physical and biochemical cellular behaviors, cell proliferation, survival, migration, and invasion (43). This pathway is also related to insulin signaling, glycogen synthesis, and differentiation in muscle cells. Some studies have shown that PA promotes the architectural remodeling of skeletal muscle in response to mechanical stimuli, such as what occurs during physical exercise (44–47). In addition, the PI3/Akt pathway is together with this pathway in the regulation against insulin resistance, being pathways that act in the protection against type 2 diabetes mellitus (48,49).

It is possible to notice that all these pathways are interconnected with the PI3K/Akt pathway, the pathway that showed the lowest FDR value. However, nothing in the literature relates all these pathways to lean mass or glucose metabolism. In parallel, the PI3k/Akt pathway demonstrated a large difference in FDR value compared to the other pathways (approximately 10^-1^ lower). This could indicate that, as it was more affected, its relationship with the other pathways may have influenced the difference in the methylation profile of these pathways between the LLM and MLM groups.

Activation of the PI3K pathway can stimulate musculoskeletal hypertrophy. The effects of this pathway occur to a considerable extent when they are in parallel with insulin-like growth factor 1 (IGF-1) signaling. One of the main activities of IGF-1 in hypertrophy is the ability to activate the Phosphoinositide 3-kinase (PI3K)/Akt signaling pathway. Serine-threonine protein kinase (Akt) has the ability to induce protein synthesis and block transcriptional upregulation of critical mediators of muscle atrophy (ubiquitin ligases E3, MuRF1, MAFbx), phosphorylating and inhibiting the nuclear translocation of the FOXO family. Of transcription factors (50–58).

Furthermore, signaling between IGF1 and PI3K/Akt may inhibit the effects of myostatin. This protein inhibits myoblast differentiation, blocks the Akt pathway, and deregulates the Akt/mTOR/p70S6 protein synthesis pathway. Thus, myostatin inhibition can cause an increase in skeletal muscle size, stimulating protein differentiation and synthesis (50,51).

The PI3K/Akt pathway is also related to glucose and insulin metabolism. This PICK1-activated pathway results in increased GLUT2 expression, functionally protects pancreatic β cells, promotes insulin secretion, and slows the progression of type 2 diabetes mellitus (59). PI3K/Akt also contributes to protection against neural apoptosis and diabetic encephalopathy, which is stimulated by glucose fluctuation (60). Ligands such as leptin, insulin, GLP, and growth factors act in the PI3K/Akt pathway and have essential roles in the functioning of this pathway. Among these ligands, insulin is the primary regulator of this pathway. Insulin, when it is secreted after a meal, promotes an increase in glucose utilization and a reduction in gluconeogenesis in the liver and muscles; increased deposition of lipids in the body; reduced circulation of free fatty acids in adipose tissue; reduced appetite in the brain; and increased production of insulin in the pancreas; as well as regulating the balance of lipid and glucose metabolism by stimulating the functioning of the PI3K/Akt pathway. However, excess energy, as in obesity, can impair the signaling of this pathway, causing insulin resistance (61).

From the results of our work, it is not possible to state that there was a difference in the expression of genes. However, the LLM group showed hypermethylation of the genes of the PI3K/Akt pathway, which suggests impairment in glucose metabolism, and body composition (with a lower chance of presenting a higher percentage of lean mass and a greater chance of presenting a higher percentage of body fat), and muscle functionality (7,8,50–61). However, in our study, there was no statistically significant difference between groups in blood tests of glucose metabolism and muscle functionality (p < 0.05) - with longer follow-up in future studies, any differences may become apparent. This indicates that the difference in body composition of lean mass was the determining factor in differentiating the methylation profile of the genes of the PI3K/Akt pathway between the groups. In addition, this exploratory study points to the need for cause and effect studies that seek to analyze potential associations of the aforementioned pathways with the PI3K/Akt pathway and the control of muscle trophism.

The DMR analysis detected five regions of specific genes that presented sites with statistical differences between the groups for M-values. The areas of these genes had a different methylation profiles according to the group (LLM and MLM). The expression of the protocadherin gamma gene group (PCDHGA4 and PCDHGA2) is negatively correlated with muscle strength. In addition, it may be related to muscle denervation and reinnervation (62). Although no studies demonstrate the relationship between the expression of PRSS22, C170rf97, and HLA-DPB15 genes and lean mass, studies show variations in the human leukocyte antigen (HLA) gene polymorphism are related to sarcopenia. In parallel, the expression of these genes is associated with impaired glucose metabolism (63,64).

In summary, our study demonstrated significant differences in methylation profiles between women aged 50 to 70 years who have a higher percentage of lean mass in relation to those with a lower percentage of lean mass in the context of aging. The PI3K/Akt pathway showed the lowest FDR value, and all other pathways are related to it, suggesting that this pathway may be the most affected by lean mass composition. In contrast, it was impossible to analyze genes’ expression through mRNA. Thus, it is not possible to state whether these differences in the methylation profile alter the expression level of genes that were methylated differently. In addition, macronutrients ingested in the diet were not measured. There, it is also not possible to know whether there was a difference in the intake of proteins, carbohydrates, and lipids between the groups and if this could influence the percentage of lean mass and the methylation profile of the participants. Our results demonstrate a differentiation between specific sites of different genes, which have essential functions in body composition and energy metabolism, supporting future studies that aim to relate lean mass with epigenetics. Therefore, it is suggested that future studies analyze the relationship between lean mass and gene expression through mRNA for a better understanding of the epigenetic influence on the control of body composition and validation of epigenetic biomarkers to predict reduced levels of muscle mass.

